# Humoral correlates of protection against *Mycobacterium tuberculosis* following intravenous Bacille Calmette-Guérin vaccination in rhesus macaques

**DOI:** 10.1101/2023.07.31.551245

**Authors:** Edward B. Irvine, Patricia A. Darrah, Shu Wang, Chuangqi Wang, Ryan P. McNamara, Mario Roederer, Robert A. Seder, Douglas A. Lauffenburger, JoAnne L. Flynn, Sarah M. Fortune, Galit Alter

## Abstract

Altering the route of Bacille Calmette-Guérin (BCG) immunization from low-dose intradermal vaccination to high-dose intravenous (IV) vaccination resulted in a high level of protection against *Mycobacterium tuberculosis* (*Mtb*) infection, providing an opportunity to uncover immune correlates and mechanisms of protection. In addition to strong T cell immunity, IV BCG vaccination was associated with a robust expansion of humoral immune responses that tracked with bacterial control. However, given the near complete protection afforded by high-dose IV BCG immunization, a precise correlate of immune protection was difficult to define. Here we leveraged plasma and bronchoalveolar lavage fluid (BAL) from a cohort of rhesus macaques that received decreasing doses of IV BCG and aimed to define the correlates of immunity across macaques that experienced immune protection or breakthrough infection following *Mtb* challenge. We show an IV BCG dose-dependent induction of mycobacterial-specific humoral immune responses, both in the plasma and in the airways. Moreover, antibody responses at peak immunogenicity significantly predicted bacterial control following challenge. Multivariate analyses revealed antibody-mediated complement and NK cell activating humoral networks as key functional signatures associated with protective immunity. Collectively, this work extends our understanding of humoral biomarkers and potential mechanisms of IV BCG mediated protection against *Mtb*.

## INTRODUCTION

*Mycobacterium tuberculosis* (*Mtb*), the causative agent of tuberculosis (TB), caused an estimated 1.6 million deaths in 2021 [1]. Antibiotics to treat TB exist, yet their effectiveness is hampered by imperfect patient adherence to lengthy drug regimens and emerging antibiotic resistance [1]. Like other infectious diseases, mass vaccination against TB would provide the most efficient intervention to control the epidemic. However, Bacille Calmette-Guérin (BCG), the only approved TB vaccine, fails to effectively prevent *Mtb* infection or disease in adults [2]. In the absence of a defined correlate of protection against *Mtb*, the design of next-generation vaccines has proven difficult.

While standard intradermal (ID) BCG administration results in limited protection against pulmonary disease in adolescents/adults, recent data demonstrate that simply switching the route of BCG administration can lead to profound protection against *Mtb* infection [3]. Specifically, while ID immunization of rhesus macaques with BCG lead to *Mtb* infection and disease in all macaques, macaques that received a high dose (5e+07 colony forming units (CFUs)) of intravenous (IV) BCG exhibited a remarkable level of protection from *Mtb* infection, with 6 out of 10 macaques exhibiting no detectable *Mtb* burden and 9 out of 10 harboring less than 100 CFU at the time of necropsy [3]. Although the mechanisms of protection are yet to be strictly defined, IV BCG drove the persistent activation of lung-localized myeloid cells [4], and induced a broad expansion in polyfunctional T cells and antibody titers in the periphery and lungs compared to ID BCG [3,5]. Furthermore, antigen-specific IgM responses were among the strongest humoral signatures of IV BCG induced protection [5]. However, due to the limited number of IV BCG vaccinated rhesus macaques with breakthrough infections in the original study [3], precise correlates of IV BCG mediated protection were unable to be defined.

To define IV BCG induced correlates of immunity, a follow-up dose de-escalation study was performed, in which rhesus macaques were vaccinated with decreasing doses of IV BCG prior to *Mtb* challenge [6]. Computational analysis of immune data identified CD4 T cell functions and NK cell numbers to be dose-independent primary correlates of IV BCG-mediated protection [6]. While simple LAM- and PPD-specific antibody titer measurements were captured in the original immune correlates analysis [6], we hypothesized that a more extensive survey of antibody responses elicited by IV BCG immunization may unveil novel correlates of protection and provide unique insight into potential humoral mechanisms of *Mtb* control. Detailed antibody profiling of plasma and BAL collected over the course of this vaccination study revealed an IV BCG dose-dependent induction of functional mycobacterial-specific humoral immunity. Further, antibody profiles in the plasma and BAL at peak immunogenicity significantly predicted *Mtb* infection outcome. Finally, multivariate linear modeling of longitudinal antibody data identified antigen-specific IgM responses, along with antibody complement and NK cell activation as key surrogates of *Mtb* control. These data point to novel biomarkers associated with highly protective immunity to *Mtb*, and provide insights into the mechanisms through which antibodies may drive protection within the lung.

## RESULTS

### Dose dependent induction of systemic humoral immune responses by IV BCG

IV BCG provides striking protection against *Mtb* in rhesus macaques [3]. To identify the correlates of protection, a vaccine dose de-escalation study was designed to elicit a continuum of IV BCG induced immunity and potentially a range of protective outcomes after challenge [6]. Specifically, rhesus macaques were immunized with decreasing doses of IV BCG ranging from 2.49e+07 to 3.88e+04 colony forming units (CFU) (**Figure 1A** and **Table S1**) [6]. Plasma and BAL samples were collected from each macaque prior to IV BCG vaccination, and at 4, 8/9, 12, and 22 weeks following vaccination (**Figure 1A**) [6]. Each macaque was challenged with *Mtb* 24 weeks following immunization (**Figure 1A**) [6]. While BAL was not collected after *Mtb* challenge to avoid perturbing the course of infection, post-challenge plasma was collected 26, 28, and 36 weeks following IV BCG vaccination (**Figure 1A**) [6]. Finally, the total thoracic CFU present in each macaque was quantified at necropsy performed 36 weeks following vaccination (12 weeks following *Mtb* challenge) (**Figures 1A,B**) [6]. As expected, the dose of IV BCG administered significantly negatively correlated with total CFU (spearman ρ: -0.414, p-value: 0.015), indicating that macaques immunized intravenously with higher doses of BCG achieved better protection from infection (**Figure 1B**). A proportion of macaques in each IV BCG dose group were protected, but several cases of breakthrough infections emerged across the spectrum of dose groups (**Figure 1B**) [6]. Overall, the cohort was divided evenly between sterilely protected macaques (n=17) and those with breakthrough infections (n=17) (**Figure 1A** and **Table S1**) [6], providing the critical heterogeneity in outcome needed to mine for humoral correlates of IV BCG mediated protection.

**Figure 1.**
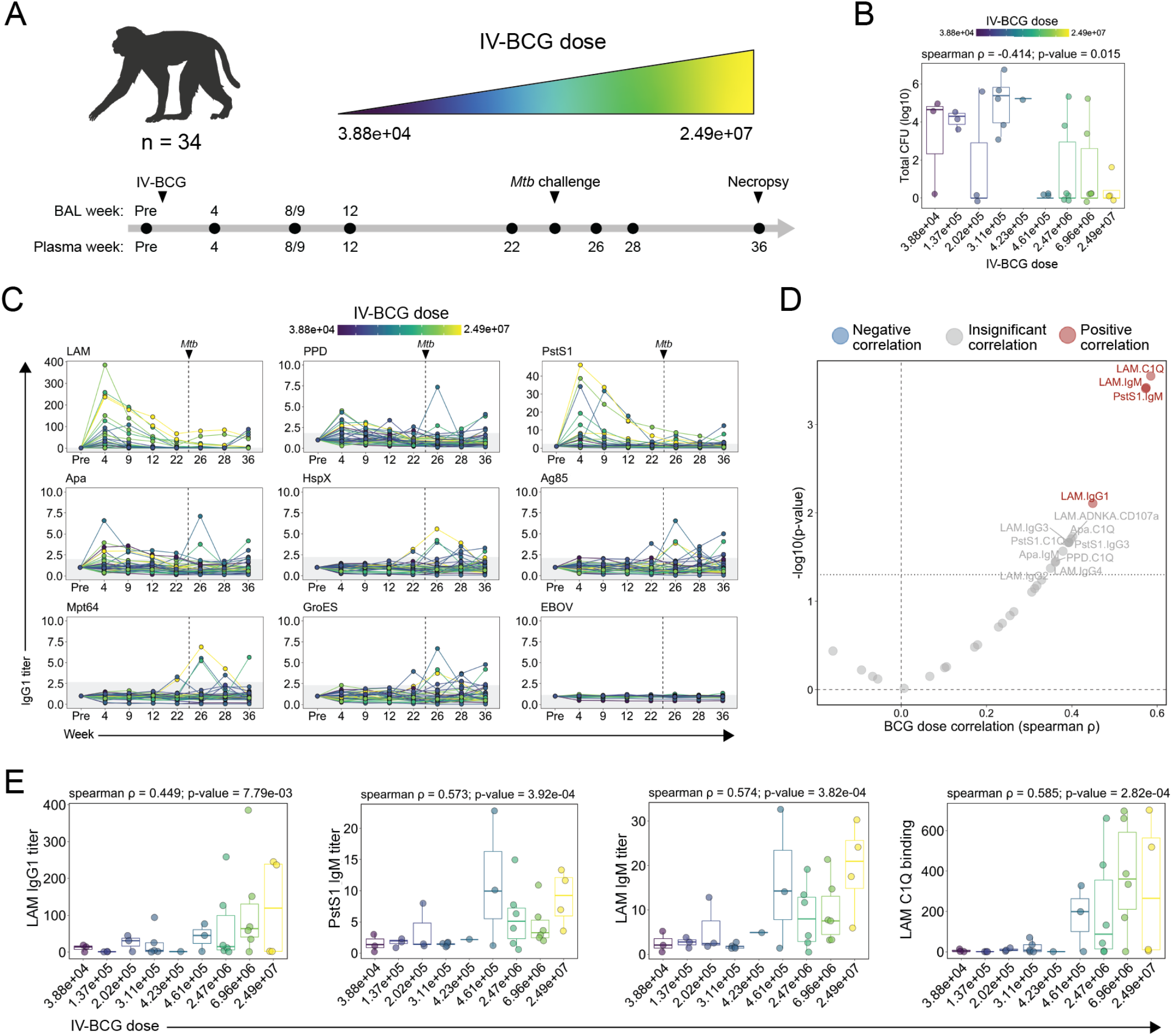
Study design and impact of IV BCG dose on plasma antibody responses. (**A**) 34 rhesus macaques were vaccinated with IV BCG at doses ranging from 3.88 × 10^4^ to 2.49 × 10^7^ CFU. Macaques were challenged with 4-17 CFU of *Mtb Erdman* 24 weeks following vaccination. Plasma and bronchoalveolar lavage fluid (BAL) was collected at multiple timepoints during the vaccination and infection phase for analysis. Week 8/9 indicates that samples were collected either 8 or 9 weeks following vaccination. Week 9 is used in future panels and figures for simplicity. (**B**) Total *Mtb* CFU measured at necropsy in each macaque [6]. Spearman correlation between log_10_(total *Mtb* CFU) and IV BCG dose indicated. (**C**) Fold change in IgG1 titers to various antigens in the plasma following IV BCG vaccination. Fold changes were calculated as fold change in Luminex median fluorescence intensity (MFI) over the pre-vaccination level for each macaque. Macaques are colored by IV BCG dose and each point represents the duplicate average from a single macaque. Dashed vertical line indicates the time of *Mtb* challenge. Grey shaded area is the background level set to 1 standard deviation above the mean MFI of the pre-vaccination samples. (**D**) Spearman correlations between IV BCG dose and each plasma antibody measurement collected at peak immunogenicity (week 4). Antibody features with low signal (average fold change less than 1.25) were removed prior to IV BCG dose correlation analysis. Black dotted horizontal line indicates unadjusted p-value of 0.05. Red and blue dots represent antibody features with a significant positive, or negative correlation with IV BCG dose respectively following multiple testing correction (Benjamini–Hochberg adjusted p-value < 0.05) [16]. (**E**) Spearman correlations between selected plasma antibody features at peak immunogenicity (week 4) and IV BCG dose.

We first queried the extent to which plasma mycobacterial-specific antibody titers, Fc*γ*-receptor binding, and antibody effector functions tracked with IV BCG dose. Systems serology, an agnostic antibody profiling platform [7], was used to probe IV BCG induced mycobacterial-specific antibody profiles across a panel of 9 mycobacterial-antigens with distinct biology and molecular function. In addition to purified protein derivative (PPD) which is a complex mixture of mycobacterial antigens [8], mycobacterial-specific antibodies were probed against secreted proteins (Ag85 and Mpt64) [9,10], cell-membrane proteins (PstS1 and Apa) [11,12], a cell wall glycolipid (lipoarabinomannan (LAM)) [13], intracellular proteins (HspX and GroES) [14,15], and an Ebola virus negative control protein (EBOV). Antibody titers over time were assessed by computing the fold change in Luminex MFI over the pre-vaccination level for each macaque at each timepoint.

Plasma IgG1, IgA and IgM antibody titers commonly peaked 4 weeks following vaccination, then waned over time (**Figure 1C** and **S1A,B**). IgG1 titers specific to LAM, PstS1, Apa, and PPD were induced by vaccination, whereas negligible vaccine-induced IgG1 titers were observed to HspX, Ag85, Mpt64, GroES, or the EBOV negative control antigen in the plasma (**Figure 1C**). IV BCG induced IgA and IgM responses were largely limited to LAM and PstS1 (**Figure S1A,B**). Furthermore, several macaques mounted antibody responses to HspX, Ag85, Mpt64, and GroES after challenge (**Figure 1C** and **S1A,B**), pointing to distinct antibody reactivity patterns induced by BCG vaccination and *Mtb* infection.

We next integrated the 90 plasma antibody features evaluated to define those most heavily influenced by vaccine dose at peak immunogenicity (week 4). Antibody features with low signal (average fold change less than 1.25) were first removed, then the remaining features with detectable vaccine-induced signal at week 4 were each individually correlated with BCG dose (**Figure 1D**). LAM-specific C1Q (spearman ρ: 0.585, p-value: 2.82e-04), LAM-specific IgM (spearman ρ: 0.574, p-value: 3.82e-04), LAM-specific IgG1 (spearman ρ: 0.449, p-value: 7.79e-03), and PstS1-specific IgM (spearman ρ: 0.573, p-value: 3.92e-04) were the only antibody features at peak immunogenicity that remained significantly positively correlated with BCG dose after multiple testing correction (**Figure 1D,E**) [16]. Other features including LAM-specific antibody-dependent NK degranulation (spearman ρ: 0.402, p-value: 0.018) and Apa-specific IgM (spearman ρ: 0.379, p-value: 0.027) and C1Q binding antibodies (spearman ρ: 0.398, p-value: 0.020) also positively correlated with IV BCG dose, but these features did not remain significant following multiple testing correction (**Figure 1D**). Together, these data indicate that IV BCG drives a dose-dependent change in mycobacterial-specific peripheral humoral immunity, with antigen-specific IgM and C1Q binding responses tracking particularly closely with the dose of vaccine administered.

### Dose-dependent induction of airway humoral immune responses by IV BCG

We next evaluated mycobacterial-specific antibody titers, Fcγ-receptor binding, and antibody effector activity in the bronchoalveolar lavage fluid (BAL). Unlike plasma samples, vaccine-induced antibody responses were detected to all mycobacterial antigens, at least in a subset of macaques (**Figure 2A** and **S2A,B**). IV BCG drove particularly robust LAM-specific responses in the airways, reaching up to a 600-fold increase in LAM-specific IgG1 titers in some macaques at week 4 (**Figure 2A** and **S2A,B**). Like plasma antibody profiles, BAL IgG1, IgA, and IgM titers peaked at week 4, and then waned over time (**Figure 2A** and **S2A,B**). Yet even 12 weeks following IV BCG immunization, the closest timepoint available before *Mtb* challenge, several macaques in the higher dosage groups maintained antibody titers above the pre-vaccination level (**Figure 2A** and **S2A,B**). As expected, neither IgG1 nor IgA responses reactive to the EBOV negative control protein were observed at any timepoint (**Figure 2A** and **S2A,B**), although a low level of IgM reactivity was observed likely due to the polyreactive nature of pentameric IgM (**S2B**) [17].

**Figure 2.**
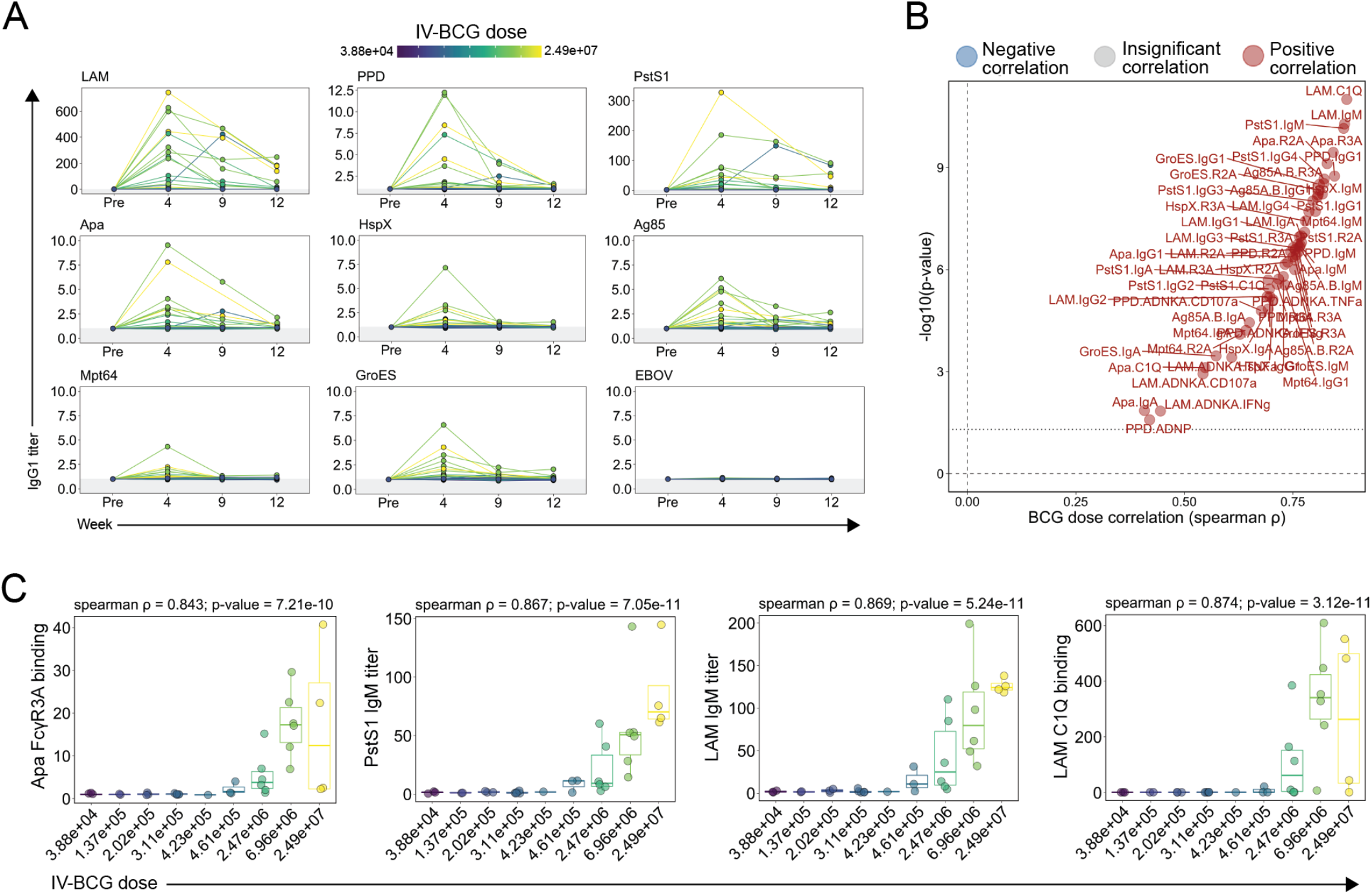
Impact of IV BCG dose on BAL antibody responses. (**A**) Fold change in IgG1 titers to various antigens in the BAL following IV BCG vaccination. Fold changes were calculated as fold change in Luminex median fluorescence intensity (MFI) over the pre-vaccination level for each macaque. Macaques are colored by IV BCG dose and each point represents the duplicate average from a single macaque. Grey shaded area is the background level set to 1 standard deviation above the mean MFI of the pre-vaccination samples. (**B**) Spearman correlations between IV BCG dose and each plasma antibody measurement collected at peak immunogenicity (week 4). Antibody features with low signal (average fold change less than 1.25) were removed prior to IV BCG dose correlation analysis. Black dotted horizontal line indicates unadjusted p-value of 0.05. Red and blue dots represent antibody features with a significant positive, or negative correlation with IV BCG dose respectively following multiple testing correction (Benjamini–Hochberg adjusted p-value < 0.05) [16]. (**C**) Spearman correlations between selected BAL antibody features at peak immunogenicity (week 4) and IV BCG dose.

We next integrated the 90 BAL antibody measurements captured at peak immunogenicity (week 4), removed those with low signal, and correlated the remaining antibody features with IV BCG dose. Every antibody feature with detectable vaccine-induced signal exhibited a significant positive correlation with IV BCG dose following multiple testing correction (**Figure 2B**) [16]. These significant BCG-dose correlates included antibody features across isotypes, antigens, and assays (**Figure 2B**). LAM-specific C1Q binding antibodies (spearman ρ: 0.874, p-value: 9.90e-12), Apa-specific Fc*γ*R3A binding antibodies (spearman ρ: 0.843, p-value: 3.59e-10), and LAM- (spearman ρ: 0.869, p-value: 5.24e-11) and PstS1-specific (spearman ρ: 0.867, p-value: 7.05e-11) IgM titers each correlated with IV BCG dose particularly closely (**Figure 2B,C**). These data illustrate that antibody responses that emerge in the BAL following IV BCG immunization are highly dependent on the dose of IV BCG administered.

### Early IV BCG induced antibody responses predict *Mtb* infection outcome

Antibodies represent the correlate of protection following nearly all clinically approved vaccines [18]. Hence, we next sought to determine whether peak mycobacterial-specific humoral immunity could similarly predict IV BCG induced protection. We first examined the relationship between BAL derived antibody measurements at week 4 and bacterial burden quantified at necropsy (**Figure 3A**). After multiple testing correction, a number of peak antibody features displayed a significant negative correlation with total CFU (**Figure 3A**). Specifically, PPD-specific FcγR2A (spearman ρ: -0.532, p-value: 8.91e-04) and FcγR3A (spearman ρ: -0.538, p-value: 9.06e-04) binding antibody levels were among the strongest correlates of protection, as were LAM-specific IgG2 titers (spearman ρ: -0.524, p-value: 1.20e-03) and IgM titers specific to PstS1 (spearman ρ: -0.497, p-value: 3.28e-03) and GroES (spearman ρ: -0.511, p-value: 2.37e-03) (**Figure 3A**).

**Figure 3.**
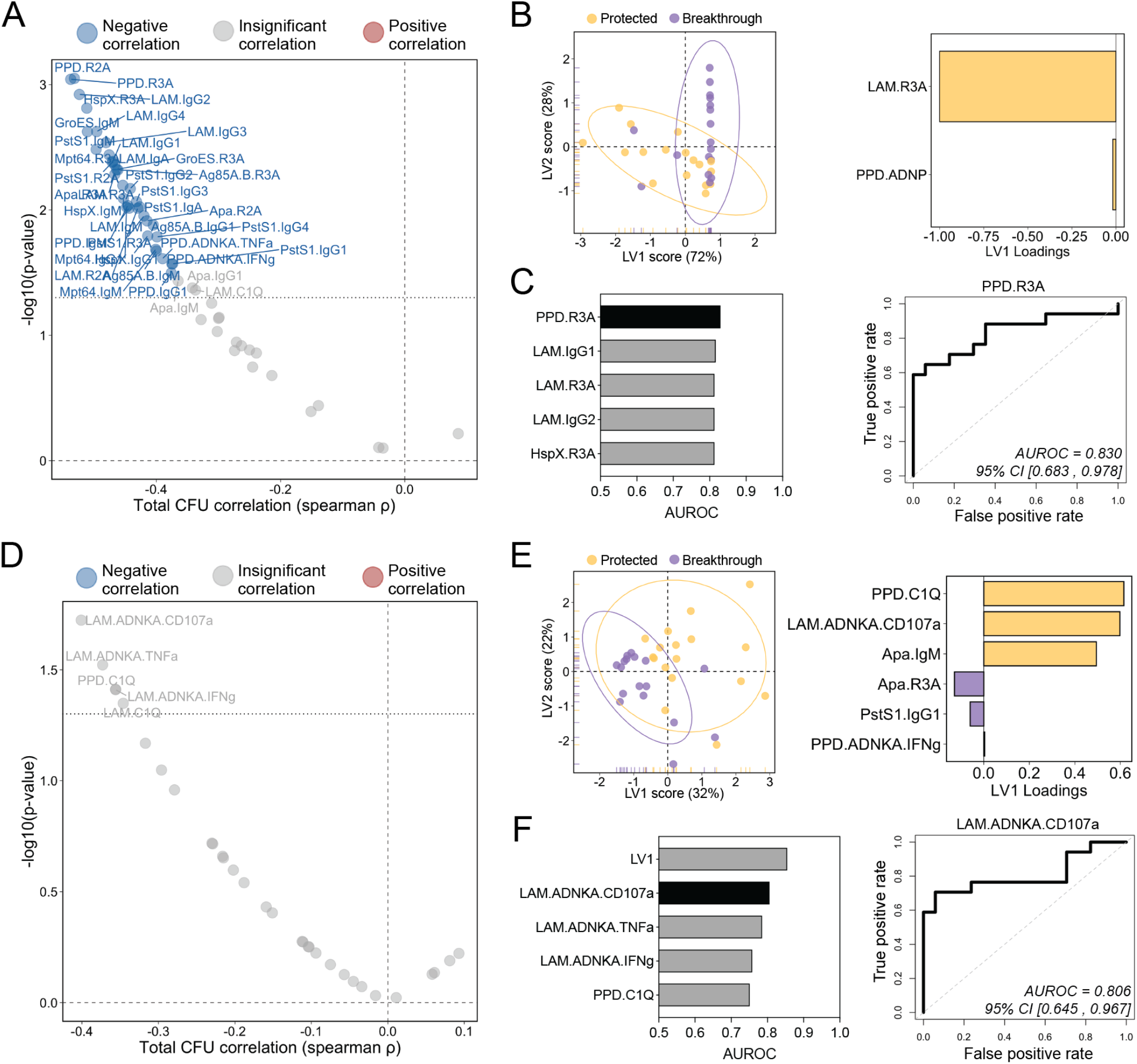
Early BAL and plasma antibody features predict *Mtb* control at necropsy. (**A** and **D**) Spearman correlations between total *Mtb* CFU measured at necropsy and each (**A**) BAL, and (**D**) plasma antibody measurement collected at peak immunogenicity (week 4). Antibody features with low signal (average fold change less than 1.25) were removed prior to correlation analysis. Black dotted horizontal line indicates unadjusted p-value of 0.05. Red and blue dots represent antibody features with a significant positive, or negative correlation with total *Mtb* CFU respectively following multiple testing correction (Benjamini–Hochberg adjusted p-value < 0.05) [16]. (**B** and **E**) PLS-DA model fit using LASSO-selected antibody features in the (**B**) BAL, and (**E**) plasma. Left: plot of latent variable (LV) 1 and 2. Ellipses represent 95-percentile normal level sets. Right: LV1 loadings. (**C** and **F**) Area under the receiver operating characteristic curve (AUROC) analysis using individual antibody features in the (**C**) BAL, and (**F**), plasma. Left: AUROC of the five most predictive features. Top individual antibody predictor in black, remaining features in grey. Right: Receiver operating characteristic plot of the top individual antibody predictor in the (**C**) BAL, and (**F**) plasma. 95% confidence intervals of the AUROC are shown in brackets.

We next sought to define a minimal mycobacterial-specific antibody signature in the BAL, captured 4 weeks following immunization, that could accurately predict IV BCG induced protection against *Mtb*. IV BCG vaccinated macaques were split into two groups to generate a sterilizing *Mtb* control outcome variable distinguishing macaques that that exhibited sterilizing protection against *Mtb* (total CFU=0; n=17), and those that experienced breakthrough infections (total CFU>0; n=17). Next, the BAL antibody profiling data were integrated, and a combination of least absolute shrinkage and selection operator (LASSO) regularization and partial least-squares discriminant analysis (PLS-DA) was implemented to identify a minimal set of features able to accurately separate the protected macaques from those with breakthrough infections (**Figure 3B**). Robust separation was observed between the two groups simply based on variation along latent variable 1 (LV1) (**Figure 3B, left**). Strikingly, as few as 2 of the total 90 features that were captured in the BAL samples were sufficient to significantly separate the groups: LAM-specific FcγR3A binding antibodies and PPD-specific antibody-dependent neutrophil phagocytosis (ADNP) (**Figure 3B, right** and **S3A**). Given the success of this relatively simple PLS-DA model, we reasoned that individual antibody features may also have strong predictive power. Thus, we next performed area under the receiver operator curve (AUROC) analysis to determine the ability of each to predict *Mtb* infection outcome (**Figure 3C**). Several peak BAL antibody features were strongly predictive of group separation on their own (**Figure 3C**). For example, LAM-specific FcγR3A binding antibodies, a key featured identified by the LASSO model (**Figure 3B, right**), had an AUROC value of 0.814 (**Figure 3C**), pointing to its considerable predictive power. BAL PPD-specific Fc*γ*R3A binding antibodies and LAM-specific IgG1 titers were also strong predictors of protection with AUROC values of 0.830 and 0.818 respectively (**Figure 3C**). Of note, the five antibody features found to be most predictive during logistic regression analysis (**Figure 3C, right**), were also significantly higher in protected macaques compared to those with breakthrough infection by Mann-Whitney analysis (**Figure S3B**).

We next aimed to determine whether *Mtb* infection outcome following IV BCG vaccination could be predicted using peak (week 4) antibody measurements in the plasma – a compartment easier to sample and thus of high-value for correlate of protection studies. Plasma LAM-specific NK cell degranulation (spearman ρ: -0.401, p-value: 0.019) as well as PPD- (spearman ρ: -0.356, p-value: 0.039) and LAM-specific (spearman ρ: -0.346, p-value: 0.045) C1Q-binding antibodies correlated most strongly with reduced *Mtb* CFU measured at necropsy (**Figure 3D**). However, individual peak immunogenicity plasma measurements did not significantly correlate with bacterial control after multiple testing correction (**Figure 3D**) [16].

To determine whether a significant multivariate signature of protection could be defined, we next generated a LASSO PLS-DA model using the week 4 plasma antibody measurements (**Figure 3E**). Robust separation of protected and breakthrough macaques was observed (**Figure 3E, left**), with significant separation driven by only 6 of the total 90 features evaluated in the plasma samples (**Figure 3E, right** and **S3C**). Interestingly, PPD-specific C1Q binding antibodies (PPD.C1Q), LAM-specific antibody-dependent NK cell degranulation (LAM.ADNKA.CD107a), and Apa IgM titers, were all selectively enriched in the plasma of protective macaques. LV1 was a strong predictor of protection, with an AUROC of 0.855 (**Figure 3F**). Furthermore, similar to the BAL, several individual plasma features measured at peak immunogenicity exhibited substantial predictive power, such as LAM-specific antibody-dependent NK cell degranulation (LAM.ADNKA.CD107a) and activation (LAM.ADNKA.TNFa), which had AUROC values of 0.806 and 0.786 respectively (**Figure 3F**). Further, the top 5 most predictive plasma antibody features were significantly higher in protected macaques by Mann-Whitney analysis (**Figure S3D**). Taken together, these data indicate that several humoral features measured 4 weeks following IV BCG immunization correlate significantly with reduced *Mtb* burden, and further suggest that in the setting of binary classification, sterilizing protection from *Mtb* can be accurately predicted using a single BAL or plasma antibody measurement collected early after vaccination.

### Convergent humoral signatures of protection before and after challenge

While previous analyses focused on correlates of protection at peak immunogenicity, we next aimed to determine whether combining all the antibody profiling data collected across timepoints and compartments may offer unique mechanistic insights into potential humoral mechanisms of *Mtb* control. Thus, all antibody data was integrated to define a minimal cross-compartment signature associated with complete protection against *Mtb* following IV BCG vaccination. Importantly, rather than using peak immunogenicity levels, here we calculated an area under the curve value for each antibody feature to capture the overall levels of the feature up to and following *Mtb* challenge (**Figure 4A**). Thus, following feature computation, each antibody feature (e.g. Plasma LAM IgG1), was transformed into 2 integrated variables describing the longitudinal behavior of the feature during the pre-challenge phase (e.g. Plasma.Pre.LAM.IgG1) and the post-challenge phase (e.g. Plasma.Post.LAM.IgG1) of the study (**Figure 4A, top**). Similarly, area under the curve values were computed for each BAL antibody feature during the pre-challenge phase (**Figure 4A, bottom**). Then all pre- and post-*Mtb* challenge features were integrated and a LASSO PLS-DA analysis was performed on the combined dataset, to define the features that most accurately captured variation between fully protected and breakthrough macaques (**Figure 4B**).

**Figure 4.**
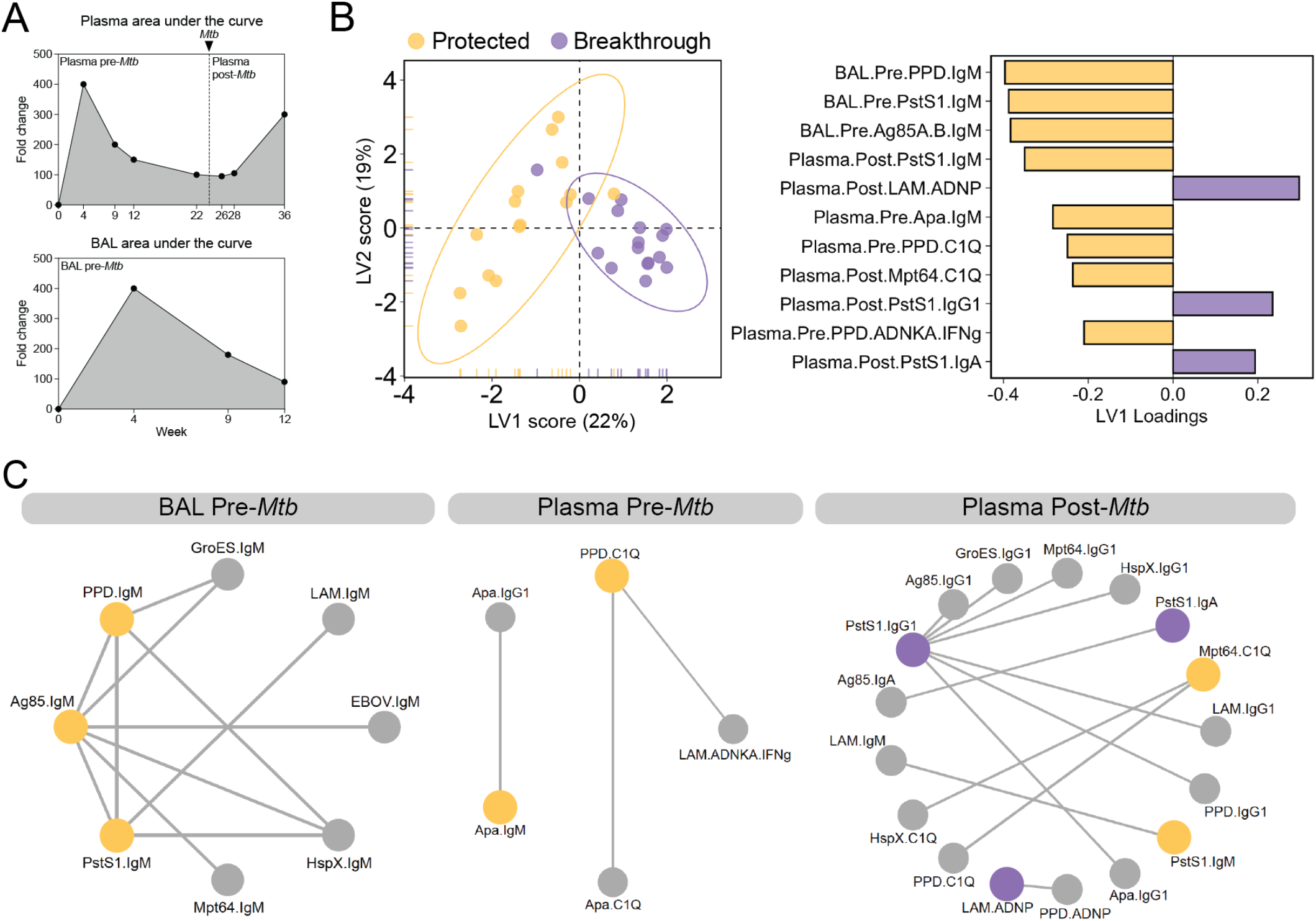
Convergent humoral signatures of protection across compartments pre- and post-*Mtb* challenge. (**A**) Fictitious data demonstrating how each antibody feature was transformed into 3 integrated variables by area under the curve computation: plasma pre-*Mtb* challenge, plasma post-*Mtb* challenge, and BAL pre-*Mtb* challenge. (**B**) PLS-DA model fit using LASSO-selected antibody features in the combined dataset. Left: plot of latent variable (LV) 1 and 2. Ellipses represent 95-percentile normal level sets. Right: LV1 loadings. (**C**) Co-correlate networks based on the pairwise correlation between the LASSO-selected antibody features, and the remaining antibody features evaluated. Networks were constructed separately for BAL pre-*Mtb* challenge features (left), plasma pre-*Mtb* challenge features (center), and plasma post-*Mtb* challenge features (right). Nodes of the LASSO-selected features enriched in protected and breakthrough macaques are yellow and purple respectively. Nodes of significant co-correlates are grey. Spearman correlations with a coefficient greater than an arbitrary threshold (pre-plasma threshold = 0.6; pre-BAL threshold = 0.9; post-plasma threshold = 0.7) and an adjusted p-value less than 0.01 after multiple testing correction are indicated by the edges [16].

Robust separation was observed across the macaque groups largely along the LV1 axis (**Figure 4B, left**). 11 of the 270 total area under the curve features in the dataset were sufficient to significantly distinguish the 2 groups, marked largely by features elevated in the protected macaques (**Figure 4B** and **S4A**). PPD-, PstS1-, and Ag85-specific IgM titers in the BAL prior to challenge were among the top features enriched among fully protected macaques (**Figure 4B, left**). Indeed, univariate analysis of these BAL LASSO-selected features found each to be significantly higher in protected macaques compared to macaques with breakthrough infection during at least one of the timepoints sampled (**Figure S4B**). In the plasma, Apa-specific IgM, PPD-specific C1Q binding antibodies, and PPD-specific NK activating antibodies were selectively enriched among protected macaques (**Figure 4B, right**). Of note, PstS1-specific IgM antibodies and Mpt64-specific C1Q binding antibodies after challenge were also enriched among protected macaques in the plasma (**Figure 4B, right**), pointing to an enrichment of IgM and complement mediated immunity in protected macaques close to the time of *Mtb* infection. Consistent with this notion, IgM titers to PstS1 and Apa were significantly higher in protected macaques compared to those with breakthrough infection at the 3 timepoints nearest to *Mtb* challenge: week 22, week 26, and week 28 (**Figure S4C**). Moreover, spearman correlations performed close to, and following *Mtb* challenge consistently identified plasma IgM and C1Q binding antibodies as significantly correlated with reduced *Mtb* burden (**Figure S4D**). By contrast, plasma LAM-specific ADNP, PstS1-IgG1, and PstS1-IgA expansion after challenge were identified by the model as linked to breakthrough infection (**Figure 4B, right**), suggesting that these responses expand selectively after infection in macaques unable to control bacterial replication.

Humoral immune responses were highly correlated likely due to the simultaneous induction of many antibody specificities after IV BCG vaccination (**Figure S5**). However, our LASSO feature selection approach selects only a subset of highly correlated features to include in the model. Therefore, to explore the broader humoral networks tracking with *Mtb* infection outcome – beyond the features selected by our model – we performed a co-correlates analysis using the selected features. Strikingly, the PPD-, PstS1-, and Ag85-specific IgM titers selected by the model as augmented in the BAL of protected macaques, correlated strongly with IgM titers to nearly all tested antigens tested, hinting at a critical role for IgM in the lung prior to *Mtb* challenge (**Figure 4C, left**). In the plasma prior to *Mtb* challenge, Apa-specific IgM titers and PPD-specific C1Q binding antibodies, which were enriched in the protected group, were strongly correlated with Apa IgG1 titers, Apa-specific C1Q binding antibodies, and LAM-specific NK cell activating antibodies (**Figure 4C, middle**). Finally in the plasma following *Mtb* challenge, Mpt64-specific C1Q binding antibodies and PstS1-specific IgM titers following infection, that were enriched in protected macaques, displayed strong correlations with additional C1Q and IgM related features including HspX- and PPD-specific C1Q binding antibodies, as well as LAM-specific IgM titers (**Figure 4C, right**), revealing a similar complement fixing IgM signature of protection prior to and following *Mtb* challenge. Conversely, LAM neutrophil phagocytosis activating antibodies together with PstS1-specific IgG1 and IgA titers enriched in macaques with breakthrough infections, strongly correlated with a myriad of additional IgG and IgA related features such as IgG1 titers across all *Mtb* antigens evaluated, and Ag85-specific IgA (**Figure 4C, right**), suggesting that IgG- and IgA-skewed responses expand selectively following breakthrough infection as a consequence of increasing bacterial antigen load. Taken together, these data indicate that IV BCG mediated protection from *Mtb* infection is linked to robust complement fixing IgM and NK cell activating antibodies.

### Humoral signatures of protection maintained after controlling for IV BCG dose

Given that IV BCG vaccination dose significantly negatively correlated with total CFU (spearman ρ: - 0.414, p-value: 0.015) (**Figure 1B**), we finally utilized logistic regression analysis to evaluate the relationship between each antibody feature and protection while controlling for IV BCG dose. Consistent with our LASSO PLS-DA modeling approach, multiple logistic regression analysis, which included IV BCG dose as a covariate, identified a complement fixing IgM and NK cell activating antibody signature of protection (**Figure 5**). Specifically, after controlling for IV BCG dose, plasma LAM-specific NK cell degranulation (LAM.ADNKA.CD107a) and activation (LAM.ADNKA.TNFa, LAM.ADNKA.IFNg) prior to *Mtb* challenge significantly associated with protection (**Figure 5**). Plasma PPD- and Apa-specific C1Q binding antibodies prior to *Mtb* challenge, and PstS1-specific IgM titers after *Mtb* challenge also remained significantly associated with protection after controlling for IV BCG dose (**Figure 5**). Conversely, BAL-specific antibody features were no longer significantly associated with protection (**Figure 5**). These findings indicate that association of mycobacterial-specific NK cell activating and C1Q binding antibody responses in the plasma with protection following IV BCG vaccination is not simply a function of vaccine dose.

**Figure 5.**
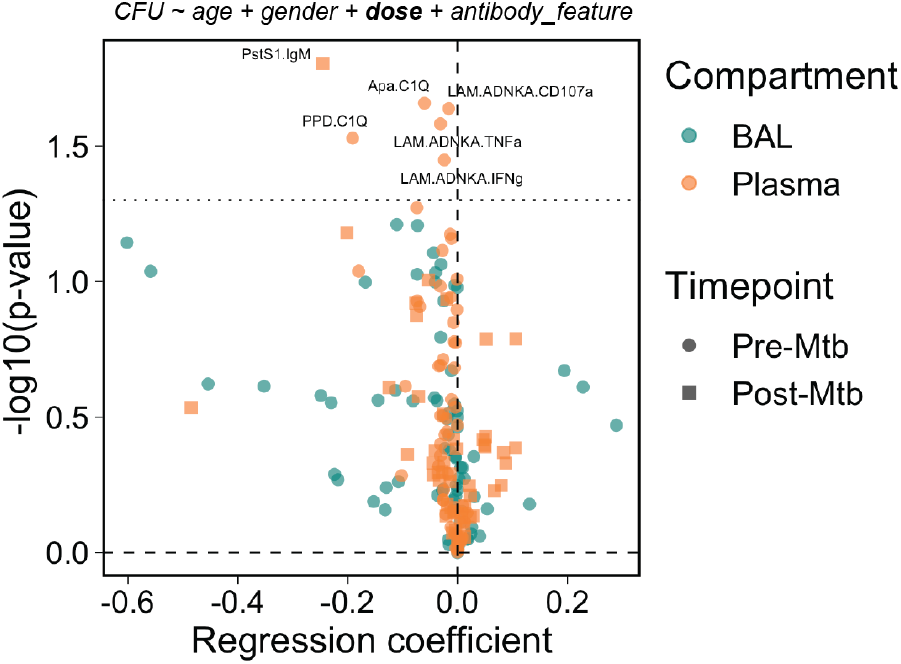
Humoral biomarkers of protection after controlling for IV BCG dose. Logistic regression to predict *Mtb* infection outcome using individual antibody features in the BAL and plasma. Each antibody feature was transformed into 3 integrated variables by area under the curve computation: plasma pre-*Mtb* challenge, plasma post-*Mtb* challenge, and BAL pre-*Mtb* challenge. Black dotted horizontal line indicates unadjusted p-value of 0.05.

## DISCUSSION

To date, IV BCG vaccination has displayed the strongest signal of protective efficacy against *Mtb* infection in non-human primates [3]. Emerging data evaluating correlates of IV BCG mediated protection indicate that a combination of systemic and respiratory compartment specific profiles are selectively augmented in protected macaques, including persistent activation in the airway myeloid cells [4], and a profound expansion of T cell and antibody immunity in the lungs and periphery [6]. While LAM- and PPD-specific antibody titer measurements were captured in the original immune correlates analysis [6], immunization with BCG likely induces a functional humoral immune response to a broad array of antigens. Thus, here we aimed to extend this analysis, to define whether a more comprehensive evaluation of antibody responses elicited by IV BCG immunization may reveal novel humoral signatures of protection.

Immune correlates play a critical role in guiding vaccine design. Recent analysis of the study presented here that included T cell responses and simple LAM- and PPD-specific antibody titer measurements identified the frequency and number of mycobacterial-specific CD4 T cells and the number of NK cells in the BAL measured over the course of the vaccination phase to be among the strongest correlates of IV BCG induced protection after controlling for IV BCG dose, while cellular immune features and antibody titers in the blood were less robust biomarkers [6]. By contrast, in the present study, humoral biomarkers associated with differential control of *Mtb* were observed in both the BAL and the peripheral circulation. Importantly, correlates: (i) linked to protective mechanisms, (ii) that are simple to measure (e.g. in the blood), and (iii) that have robust predictive power even when collected at a single cross-sectional timepoint have the greatest potential to accelerate TB vaccine design and evaluation. Hence, the top antibody predictors of IV BCG induced protection measured at peak immunogenicity (week 4), such as LAM-specific NK cell activating antibodies and C1Q binding antibodies in the plasma, may serve as easily measured markers of the quality of antigen-specific immunity induced by vaccines that may modulate *Mtb* infection. Whether the key antibody correlates of protection identified in this study are conserved across TB vaccine platforms, or instead are unique to IV BCG vaccination, remains unknown.

Our previous comparison of IV BCG-induced antibody profiles with those induced by alternate routes of BCG immunization identified robust IgM responses in the plasma and a broad expansion of humoral immunity in the BAL as markers of IV BCG induced immunity [5]. Likewise, plasma antibody features including LAM- and PstS1-specific IgM, and LAM-specific C1Q binding antibodies correlated strongly and significantly with IV BCG dose in the present study. Moreover, BAL antibody responses – across isotypes, antigens, and functions – were highly dependent on the dose of IV BCG administered, further supporting the results of the initial study in this independent vaccination cohort. While the antibody features that correlated most strongly with IV BCG dose were not exactly the same as those that correlated most strongly with protection, antigen-specific IgM/C1Q responses were also key in predicting protection prior to and following *Mtb* exposure. This consistent IgM/C1Q protective signal observed pre- and post-challenge was particularly striking and may potentially hint at a mechanistic role for complement in microbial control. C1Q, a primary marker of protective immunity identified in the present study, is the component of the classical complement pathway responsible for recognition of antibody-antigen complexes [19]. C1Q engagement of IgG or IgM on the surface of microbes precipitates the mobilization of downstream enzymatic processes that lead to the deposition of C3b on the microbial surface that can either promote opsonophagocytic clearance, or eventually form pores in the microbial membrane via the membrane attack complex [19]. While the membrane attack complex is critical in the lysis of a subset of gram-negative bacteria [19,20], it is unlikely that it can adequately penetrate the thick, complex cell envelope of mycobacteria to cause direct cell lysis [21]. Indeed, there is a paucity of evidence supporting a direct antimicrobial role for complement in *Mtb* control. Still, upon challenge, lung-localized IgM responses induced by IV BCG vaccination likely bathe the surface of *Mtb* with complement proteins, potentially altering the mechanism of bacterial uptake. Future work is required to determine whether enhanced IgM-mediated complement deposition on the surface of *Mtb* targets the bacteria for enhanced opsonophagocytic destruction, alters antigen presentation to lung-localized T cells, or simply promotes distinct, yet functionally neutral differences in immune signaling.

Previous functional comparisons pointed to enhanced NK cell activating antibodies as a key marker of protection in clinical control of *Mtb* [22,23]. Specifically, a multi-cohort analysis observed that latent tuberculosis infection was associated with higher signaling via FcγR3A, and enhanced NK cell activity [22], supporting a potential role for NK cell recruiting antibodies in the context of controlled latent TB infection. Similarly, our previous work identified an enrichment of FcγR3A-binding and NK cell-activating antibodies in controlled latent TB infection compared to individuals with active TB disease [23]. Purified IgG from latent TB patients bearing this unique Fc-effector profile also suppressed *Mtb* replication in primary macrophages more effectively than purified IgG from actively infected individuals, suggesting that these antibodies may target *Mtb*-infected macrophages [23]. Here we establish enhanced FcγR3A-binding and NK cell-activating antibodies as correlates of protection following IV BCG vaccination in rhesus macaques. Intriguingly, the present study reinforces and extends the findings from a previous multivariate analysis of the same cohort of macaques vaccinated with decreasing doses of IV BCG, which identified the number of NK cells in the BAL to be a primary correlate of protection [6]. This convergent signal across the two studies is consistent with a potential model whereby the higher availability of BCG in the lung may expand NK cells at the site of infection, while simultaneously instructing B cells to generate mycobacterial-specific antibodies able to drive enhanced NK cell activation. Hence, it is conceivable that local NK cells may be armed and directed by local antibodies to reduce *Mtb* lung burden and/or dissemination.

While correlation does not equate to causation, the correlates of protection identified herein engender mechanistic hypotheses regarding a potential role for antibodies in vaccine-induced control of *Mtb*. The persistence of these humoral signals until the time of *Mtb* challenge as well as post-challenge may suggest sustained antibody secretion from lung-localized B cells, and a potential rapid anamnestic recall of local B cell responses following *Mtb* exposure that may plausibly contribute to the protective immunity afforded by IV BCG as described for other respiratory pathogens [18,24,25]. Mechanistic studies in mice are challenging due to differences in Fc receptor expression and NK cell function compared humans [26–28]. Thus, passive transfer studies in non-human primates may be necessary to distinguish humoral correlates from mechanisms of protection, as well as to provide critical insights into how antibodies may be harnessed to drive enhanced immunity against *Mtb* more broadly. Indeed, the continued interrogation of immune correlates and mechanisms of vaccine-induced protection against *Mtb* using IV BCG as a model may inspire the formulation of safe, next-generation TB vaccination strategies which similarly elicit robust protective immunity.

## ACKNOWLEDGEMENTS

We acknowledge support from the Ragon Institute of MGH, MIT and Harvard and the SAMANA Kay MGH Research Scholar Program (to SMF and GA), the Bill and Melinda Gates Foundation (OPP1156795 to SMF and GA), and the National Institutes of Health (U54CA225088 to GA, U2CCA233262 to GA, U2CCA233280 to GA, AI150171-01 to EBI, and contract no. 75N93019C00071 to SMF, DAL, CW, JLF, and GA).

## MATERIALS AND METHODS

### Cohort information

Rhesus macaque (*Macaca mulatta*) plasma and bronchoalveolar lavage fluid (BAL) samples from the BCG dose de-escalation vaccination cohort were collected during a study performed at the Vaccine Research Center at the National Institutes of Health [6]. Macaques were challenged with *Mtb* at the University of Pittsburg and post-challenge samples as well as all data from necropsy were collected at the University of Pittsburg. Experimentation and sample collection from the study complied with ethical regulations at the respective institutions. 34 macaques were immunized with intravenous (IV) BCG at half-log increasing target doses between 4.5 and 7.5 (log_10_) colony forming units (CFUs) [6]. 24 weeks following IV BCG immunization, macaques were challenged with 4-17 CFU of *Mtb Erdman* [6]. The present cohort consists of plasma and BAL collected prior to vaccination, week 4, week 8 (Cohort B)/9 (Cohort A), week 12, and week 22 following vaccination, as well as post-*Mtb* challenge plasma samples collected at week 26, week 28, and week 36 (necropsy). BAL fluid was received as a 10X concentrate and diluted for experiments. Post-*Mtb* challenge BAL samples were not collected.

### Antigens

Panel of antigens used for antibody profiling: LAM (BEI Resources, NR-14848), PPD (Statens Serum Institute), PstS1 (BEI Resources, NR-14859), Apa (BEI Resources, NR-14862), HspX (BEI Resources, NR-14860), Ag85A and B (BEI Resources, NR-53525 and NR-53526), Mpt64 (BEI Resources, NR-49435), GroES (BEI Resources, NR-14861), and Zaire ebolavirus glycoprotein (R&D Systems, 9016-EB).

### Human research participants

Primary immune cells used in functional assays were from the blood of healthy, HIV-negative individuals. Specimens were provided coded or anonymized. Donors provided written, informed consent, and the study was approved by the institutional review board at Massachusetts General Hospital.

### Antibody levels

Antibody levels were measured as described previously [5]. In brief, protein antigens were coupled to magnetic Luminex beads by carbodiimide-NHS ester coupling [29], and LAM was coupled by DMTMM modification [30]. Coupled beads were incubated with 5μL of plasma or BAL overnight at 4°C in 384-well plates (Greiner Bio-One). The following plasma dilutions were used: IgG1 (1:30 PPD and Apa; 1:150 remaining antigens), IgG2, IgG3, and IgG4 (1:150 LAM and PstS1; 1:30 remaining antigens), IgA (1:30 all antigens), IgM (1:750 all antigens). BAL was used at 1X for all isotypes/subclasses and antigens. Beads were then washed, and mouse anti-rhesus IgG1 (clone 7H11), IgA (clone 9B9), or IgM (Life Diagnostics, clone 2C11-1-5) antibody was added at a concentration of 0.65μg/mL and incubated shaking at room temperature (RT) for 1 hour. Anti-rhesus IgG1 and IgA were from the National Institutes of Health Nonhuman Primate Reagent Resource supported by grants AI126683 and OD010976. Beads were washed, and phycoerythrin (PE)-conjugated anti-mouse IgG was added (ThermoFisher, 31861) at a concentration of 0.5μg/mL and incubated shaking at RT for 1 hour. Beads were washed, and relative antibody levels (PE MFI values) were measured using a FlexMap 3D (Luminex). Data are represented as fold-change over pre-vaccination levels. Samples were run in duplicate.

### Fcγ-receptor binding

Fcγ-receptor Luminex was performed as described previously with minor modifications [5,31]. Briefly, rhesus macaque FcγR2A and FcγR3A (acquired from the Duke Human Vaccine Institute Protein Production Facility) were biotinylated with a BirA biotin–protein ligase bulk reaction kit (Avidity). C1Q (Sigma, C1740) was biotinylated with Sulfo-NHS-LC-LC Biotin (ThermoFisher, A35358). Excess biotin was removed from each with 3-kDa-cutoff centrifugal filter units (Amicon). Antigen-coupled beads were incubated with 5μL of plasma or BAL overnight at 4°C in 384-well plates (Greiner Bio-One). The following plasma dilutions were used: FcγR2A and FcγR3A (1:150 all antigens), and C1Q (1:150 LAM; 1:30 remaining antigens). BAL was used at 1X for all isotypes/subclasses and antigens. After overnight incubation, streptavidin-PE (Agilent, PJRS25) was added to each biotinylated Fcγ-receptor in a 4:1 molar ratio and incubated rotating for 10min at RT to create the detection reagent. 500μM biotin (Avidity) was then added at a 1:100 ratio relative to the total solution volume and incubated rotating for 10min at RT. Beads were washed, and each prepared detection reagent was added at a concentration of 1μg/mL and incubated shaking for 1hr at RT. Beads were washed, and Fcγ-receptor binding (PE MFI values) was measured using a FlexMap 3D (Luminex). Data are represented as fold-change over pre-vaccination levels. Samples were run in duplicate.

### Antibody-dependent cellular phagocytosis (ADCP)

PPD and LAM ADCP assays were performed as described previously with minor changes [5]. PPD was biotinylated with Sulfo-NHS-LC-LC Biotin (ThermoFisher, A35358). LAM was biotinylated using hydrazide biotin (ThermoFisher, 21339). Excess biotin was removed from each with 3-kDa-cutoff centrifugal filter units (Amicon). Biotinylated antigen was then added to FITC-conjugated neutravidin beads (ThermoFisher, F8776) at an antigen(μg):bead(μL) ratio of 1μg:1μL for PPD, and 1μg:2μL for LAM and incubated overnight at 4°C. Excess antigen was washed away. Antigen-coated beads were then incubated with 10μL of diluted sample (plasma 1:30; BAL 1X) for 2hr at 37°C. THP-1 cells (50000 per well) were then added and incubated at 37°C for 16hr. Bead uptake was measured on a BD LSR II (BD Biosciences). Data were collected in FACSDiva (version 9.0) and analyzed using FlowJo (version 10.3). Phagocytic scores were calculated as: ((% FITC-positive cells) × (geometric mean fluorescence intensity of the FITC-positive cells)) divided by 10000. Data are represented as fold-change over pre-vaccination levels. Samples were run in duplicate.

### Antibody-dependent neutrophil phagocytosis (ADNP)

PPD and LAM ADNP assays were performed as described previously with minor changes [5,32]. Briefly, antigens were biotinylated, coupled to fluorescent neutravidin beads, and incubated with sample as described above for ADCP. During the 2hr bead and sample incubation, fresh peripheral blood from healthy donors was added at a 1:9 ratio to ACK lysis buffer (Quality Biological, 118-156-101) for 5min at RT. The blood was then centrifuged, the supernatant was removed, and leukocytes were washed with cold PBS (4°C). Leukocytes were then resuspended in R10 medium (RPMI (Sigma), 10% fetal bovine serum (Sigma), 10 mM HEPES (Corning), 2 mM l-glutamine (Corning)). After the 2hr bead and sample incubation, 50000 leukocytes were added and incubated for 1hr at 37°C. Cells were washed and stained with 1:100 diluted anti-human CD66b-Pacific Blue (BioLegend, 305112). Cells were washed and bead uptake was measured on a BD LSR II (BD Biosciences). Data were collected in FACSDiva (version 9.0) and analyzed using FlowJo (version 10.3). Phagocytic scores were calculated in the CD66b-positive cell population. Data are represented as fold-change over pre-vaccination levels. Samples were run in duplicate.

### Antibody-dependent NK cell activation (ADNKA)

PPD and LAM antibody-dependent NK cell activation was performed as described previously with minor changes [5,23]. In brief, ELISA plates (ThermoFisher, NUNC MaxiSorp flat bottom) were coated with antigen (300ng PPD or 150ng LAM) and incubated overnight at 4°C. Plates were then blocked with 5% BSA-PBS overnight at 4°C. 50μL of diluted sample (plasma 1:30; BAL 1X) was added and incubated for 2hr at 37°C. One day prior to adding the diluted sample, NK cells were isolated from healthy donors using the RosetteSep human NK cell enrichment cocktail (Stemcell) and Sepmate conical tubes (Stemcell). NK cells were incubated overnight at 1.5×10^6^ cells per mL in R10 media with 1ng/mL human recombinant IL-15 (Stemcell). After the 2hr sample incubation, the plates were washed and 50,000 NK cells, 2.5μL PE-Cy5 antihuman CD107a (BD, 555802), 0.4μL brefeldin A (5 mg/mL, Sigma) and 10μL GolgiStop (BD, 554724) were added to each well and incubated for 5hr at 37°C. Samples were then stained with 1:10 diluted PE-Cy7 anti-human CD56 (BD, 557747), APC-Cy7 anti-human CD16 (BD, 557758) and Alexa Fluor 700 anti-human CD3 (BD, 557943). Samples were washed, and then fixed using Perm A and Perm B (Invitrogen). The Perm B solution contained 1:50 diluted PE anti-human MIP-1β (BD, 550078), 1:50 diluted BV605 anti-human TNF-α (BioLegend, 502936), and 1:20 diluted FITC anti-human IFN-γ (BD, 340449). Cells were washed and fluorescence was measured on a BD LSR II (BD Biosciences). Data were collected in FACSDiva (version 9.0) and analyzed using FlowJo (version 10.3). Data are represented as fold-change over pre-vaccination levels. Samples were run in biological duplicate using NK cells from two different donors.

### AUROC analysis

The predictive power of individual antibody features was determined via area under the receiver operating characteristic curve (AUROC) analysis. 95% confidence intervals of the AUROC were computed with 10000 stratified bootstrap replicates. Antibody features with low signal (average fold change less than 1.25) were removed prior to AUROC analysis. AUROC analysis was performed using the *pROC* package [33], (version 1.18.0) in R (version 4.2.1).

### Logistic regression

Logistic regression models were generated to identify the identify the antibody features that significantly predicted protection while controlling for IV BCG dose. IV BCG dose, macaque age, and macaque gender were each included as covariates in the model in addition to the respective antibody feature. Antibody features with low AUC variance between macaques (pre-plasma AUC < 30; post-plasma AUC < 15; pre-BAL AUC < 1) were removed prior to multiple logistic regression analysis. Antibody features with an unadjusted p-value < 0.05 were considered significant markers of protection. Multiple logistic regression models were generated in R (version 4.2.1).

### LASSO PLS-DA

Least absolute shrinkage and selection operator (LASSO) regularization followed by partial least-squares discriminant analysis (PLS-DA) was performed to identify key antibody features that predict *Mtb* infection outcome as previously described with minor changes [5]. Briefly, 1000 bootstrap datasets of the systems serology data were generated. A LASSO model in which the lambda hyperparameter was chosen via fivefold cross-validation was then fit on each bootstrap dataset. Variable inclusion probabilities – the proportion of bootstrap replications in which the coefficient estimate is not zero – were computed for each antibody feature. Data were z-scored, and PLS-DA models were fit using a grid of variable inclusion probability cutoffs in a fivefold cross-validation framework repeated 1000 times. Model accuracy ((1 − balanced error rate) × 100) was computed for each variable inclusion probability cutoff during cross-validation. To visualize the models, the first and second latent variable (LV) from the optimal PLS-DA models are plotted, as are the loadings of LV1 which highlight the contribution of each feature to variation in *Mtb* infection outcome. Antibody features with low signal (average fold change less than 1.25) were removed prior to the week 4 LASSO PLS-DA. Likewise, antibody features with low AUC variance between macaques (pre-plasma AUC < 30; post-plasma AUC < 15; pre-BAL AUC < 1) were removed prior to the AUC LASSO PLS-DA. LASSO was implemented using the *glmnet* package (version 3.0-2) [34], and PLS-DA models were implemented using the *mixOmics* package (version 6.10.9) [35], in R (version 4.2.1).

Co-correlate networks were constructed based on the pairwise correlation between the LASSO-selected antibody features which distinguish *Mtb* infection outcome, and the remaining antibody features evaluated. Spearman correlations with a coefficient greater than an arbitrary threshold (pre-plasma threshold = 0.6; pre-BAL threshold = 0.9; post-plasma threshold = 0.7) and an adjusted p-value less than 0.01 after multiple testing correction are shown. Correlation networks were generated using the *NetworkX* library in Python (version 3.9.12).

### Statistics

Spearman correlations were computed in R (version 4.2.1). Adjusted p-values in the correlation volcano plots and network analyses were calculated using the Benjamini-Hochberg procedure [16].

## DATA AVAILABILITY

All data and metadata associated with this study are available in the main text, Supplementary Information and/or at https://fairdomhub.org/studies/XXXXX.

## CODE AVAILABILITY

Scripts to reproduce the computational analyses presented in the paper are available at https://github.com/eirvine94/IVBCGDD_antibody_manuscript.

## AUTHOR CONTRIBUTIONS

Conceptualization: EBI, PAD, MR, RAS, DAL, JLF, SMF, GA. Methodology: EBI, SW, CW. Software: EBI. Validation: EBI. Formal analysis: EBI. Investigation: EBI, PAD. Resources: PAD, MR, RAS, JLF, SMF, GA. Data curation: EBI, PAD. Writing original draft: EBI. Review and editing: all authors. Visualization: EBI. Supervision: MR, SMF, GA. Project administration: RPM, MR, RAS, DAL, JLF, SMF, GA. Funding acquisition: EBI, MR, RAS, DAL, JLF, SMF, GA.

