## Supplemental Information for "Humoral correlates of protection against *Mycobacterium tuberculosis* following intravenous Bacille Calmette-Guérin vaccination in rhesus macaques"

**This file includes:**

Table S1
Figure S1
Figure S2
Figure S3
Figure S4
Figure S5

| | $3.88\times10^4$ | $1.37\times10^5$ | $2.02\times10^5$ | $3.11\times10^5$ | $4.23\times10^5$ | $4.61\times10^5$ | $2.47\times10^6$ | $6.96\times10^5$ | $2.49\times10^7$ |
|---|---|---|---|---|---|---|---|---|---|
| Dose (CFU) | 3 | 3 | 3 | 5 | 1 | 3 | 6 | 6 | 4 |
| Cohort | A | B | A | B | B | A | A | A | A |

**Table S1 | IV-BCG dose de-escalation cohort.** 34 rhesus macaques across 2 cohorts (A, B) were vaccinated with intravenous IV BCG at doses ranging from $3.88\times10^4$ to $2.49\times10^7$ CFU.

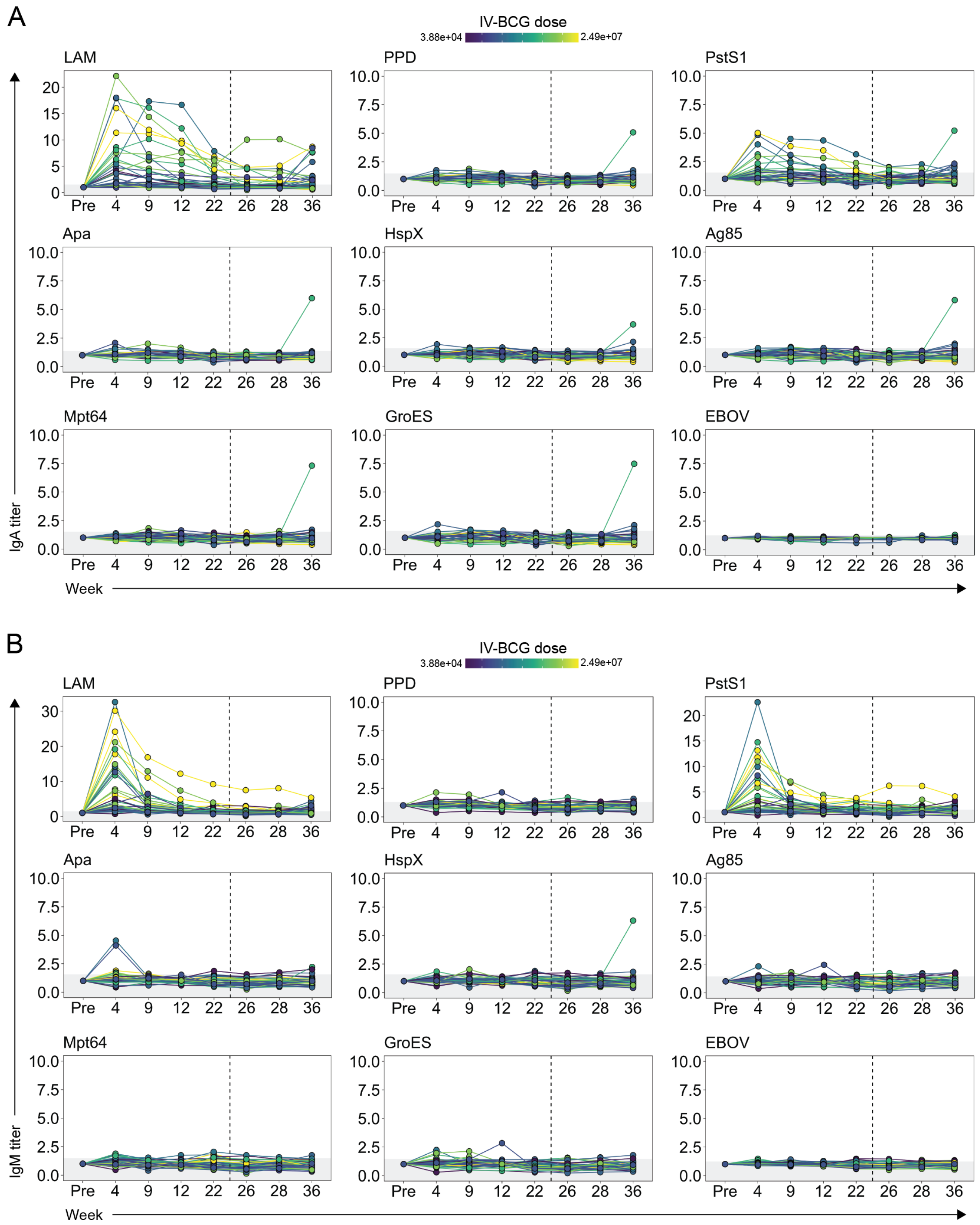


**Figure S1 | Plasma IgA and IgM titers elicited by different doses of IV-BCG immunization**. Fold change in (**A**) IgA, and (**B**) IgM titers to various antigens in the plasma following IV-BCG vaccination. Fold changes were calculated as fold change in Luminex median fluorescence intensity (MFI) over the pre-vaccination level for each macaque. Macaques are colored by IV-BCG dose and each point represents the duplicate average from a single macaque. Dashed vertical line indicates the time of *Mtb* challenge. Grey shaded area is the background level set to 1 standard deviation above the mean MFI of the pre-vaccination samples.

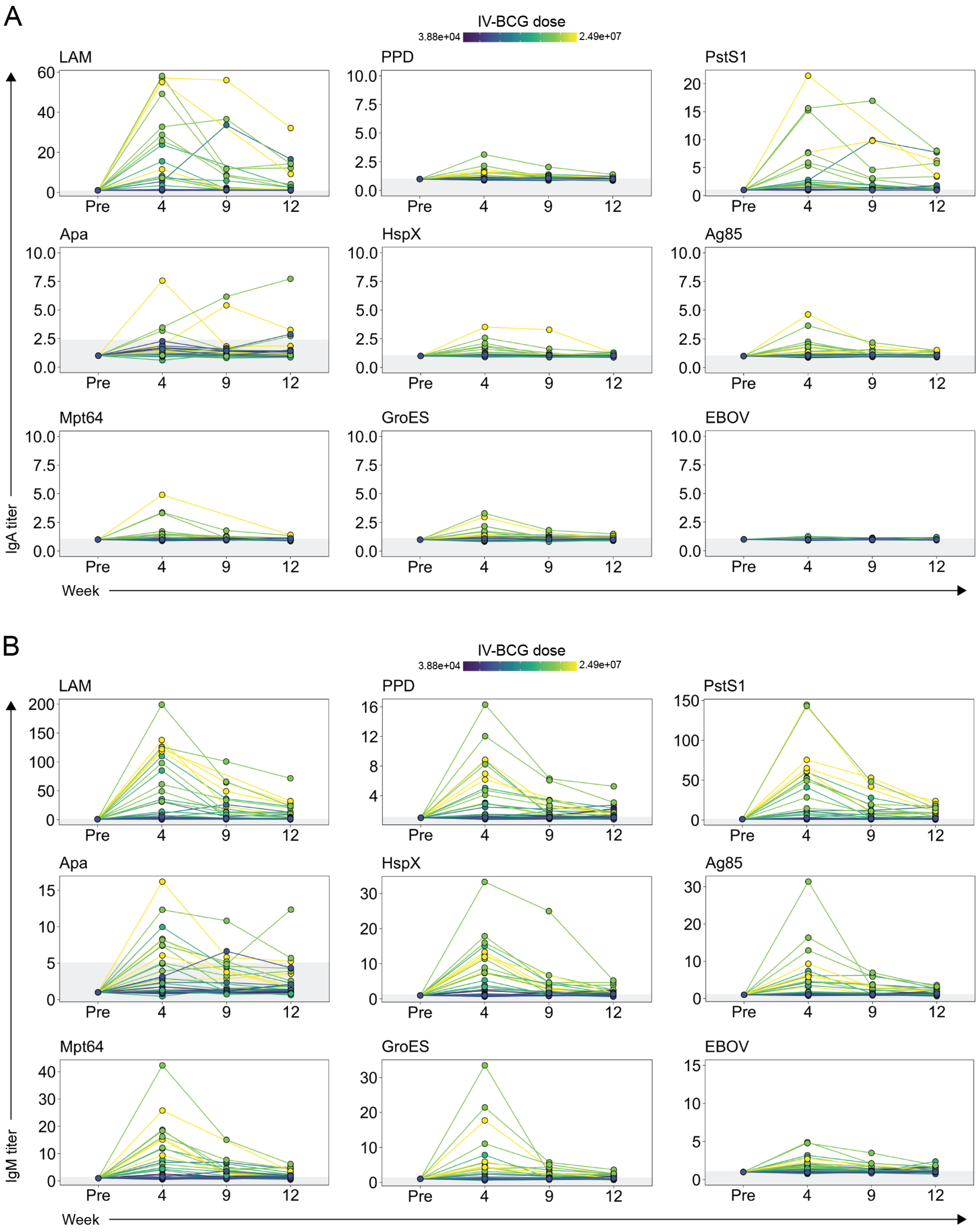


**Figure S2 | BAL IgA and IgM titers elicited by different doses of IV-BCG immunization**. Fold change in (**A**) IgA, and (**B**) IgM titers to various antigens in the BAL following IV-BCG vaccination. Fold changes were calculated as fold change in Luminex median fluorescence intensity (MFI) over the pre-vaccination level for each macaque. Macaques are colored by IV-BCG dose and each point represents the duplicate average from a single macaque. Grey shaded area is the background level set to 1 standard deviation above the mean MFI of the pre-vaccination samples.

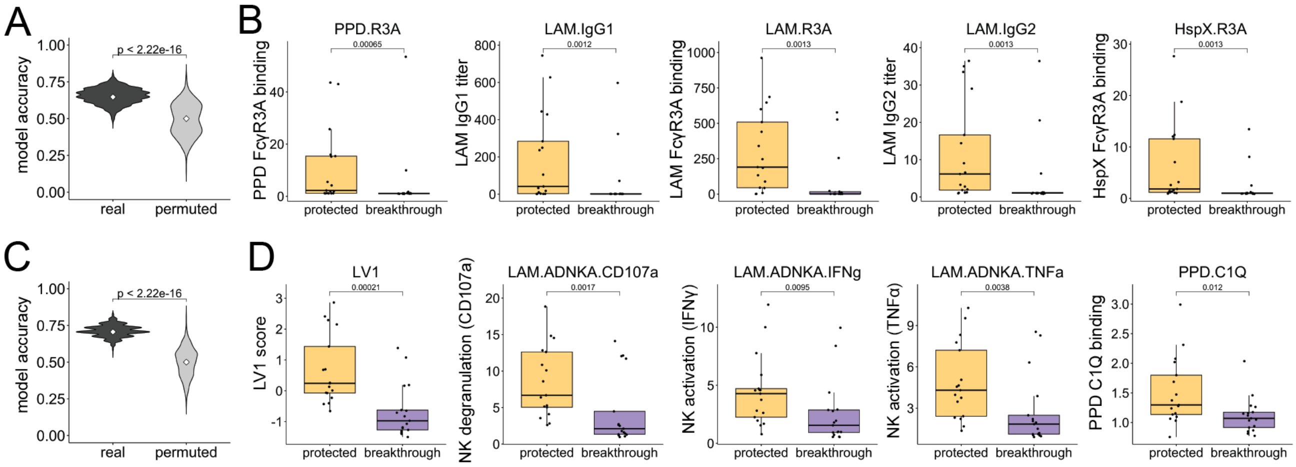


**Figure S3 | Distinguishing protected macaques from those with breakthrough infections using antibody measurements at peak immunogenicity.** (**A** and **C**) Permutation test to assess the significance of PLS-DA models fit using LASSO-selected antibody features in the (**A**) BAL, and (**C**) plasma. Balanced classification accuracy of the real model is compared with that of permuted models with the macaque outcome group labels randomly shuffled. White diamond indicates group median. Two-tailed Mann-Whitney U test. (**B** and **D**) Boxplots comparing humoral titer/activity for the top five most predictive antibody features in the (**B**) BAL, and (**D**) plasma between protected IV-BCG vaccinated macaques and those with breakthrough *Mtb* infections. Y-axes indicate fold change over the pre-vaccination level for each feature at week 4. Two-tailed Mann-Whitney U test.

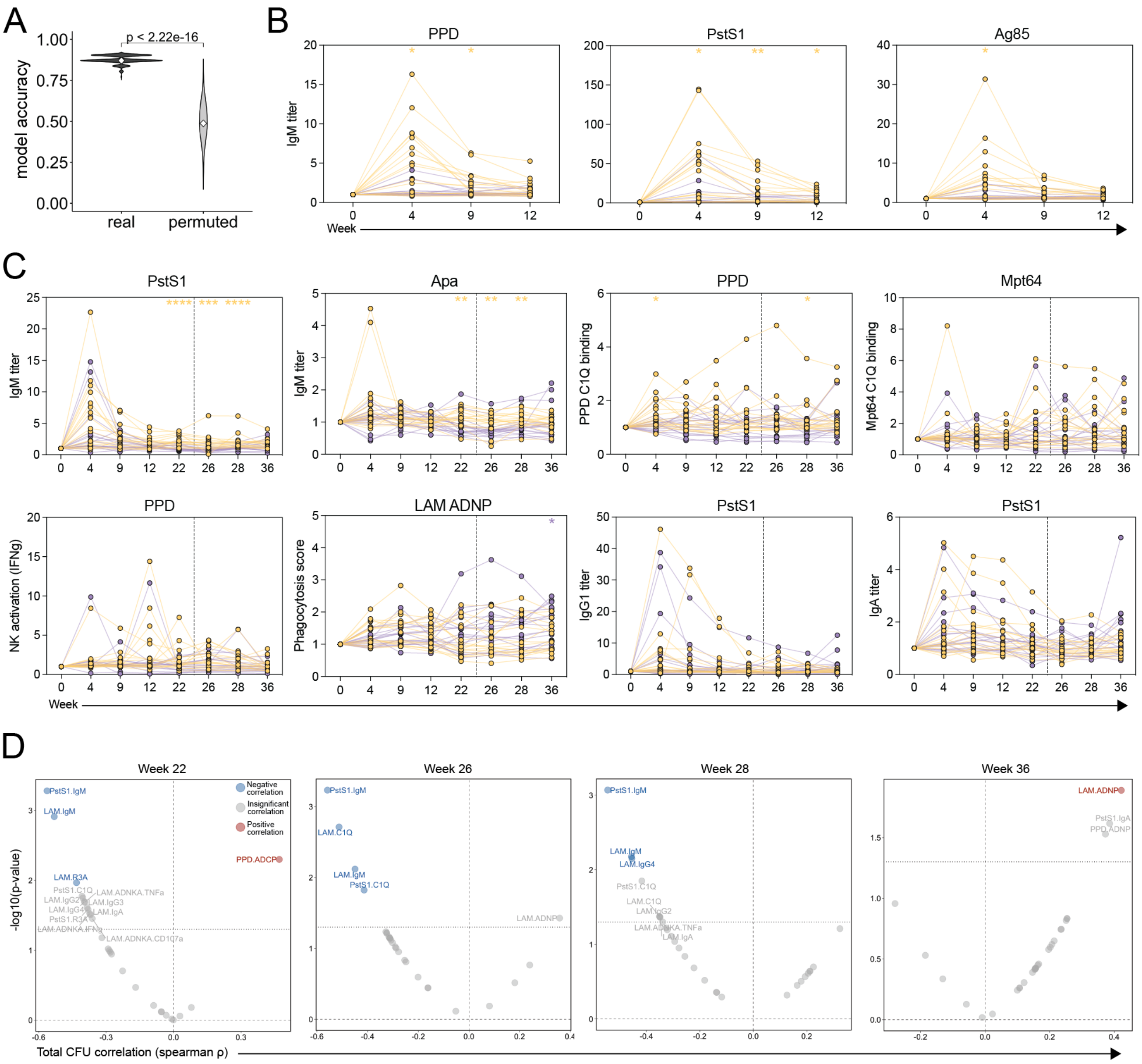


**Figure S4 | Antibody features associated with *Mtb* infection outcome in the vaccination and infection phase.** (**A**) Permutation test to assess the significance of PLS-DA model fit using LASSO-selected antibody features in the combined dataset. Balanced classification accuracy of the real model is compared with that of permuted models with the macaque outcome group labels randomly shuffled. (**B** and **C**) Longitudinal plots of the LASSO-selected features in the (**B**) BAL, and (**C**) plasma. Protected and breakthrough macaques are colored yellow and purple respectively. Fold change in each feature over the pre-vaccination level is plotted for each macaque. Two-tailed Mann-Whitney U test performed at each timepoint comparing protected and breakthrough macaques. p-value<0.05 (*), p-value<0.01 (**), p-value<0.001 (***), p-value<0.0001 (****). (**D**) Spearman correlations between total *Mtb* CFU measured at necropsy and each plasma antibody measurement collected at week 22, week 26, week 28, and week 36. Antibody features with low signal (average fold change less than 1.25) were removed prior to correlation analysis. Black dotted horizontal line indicates unadjusted p-value of 0.05. Red and blue dots represent antibody features with a significant positive, or negative correlation with total *Mtb* CFU respectively following multiple testing correction (Benjamini–Hochberg adjusted p-value < 0.05) [15].

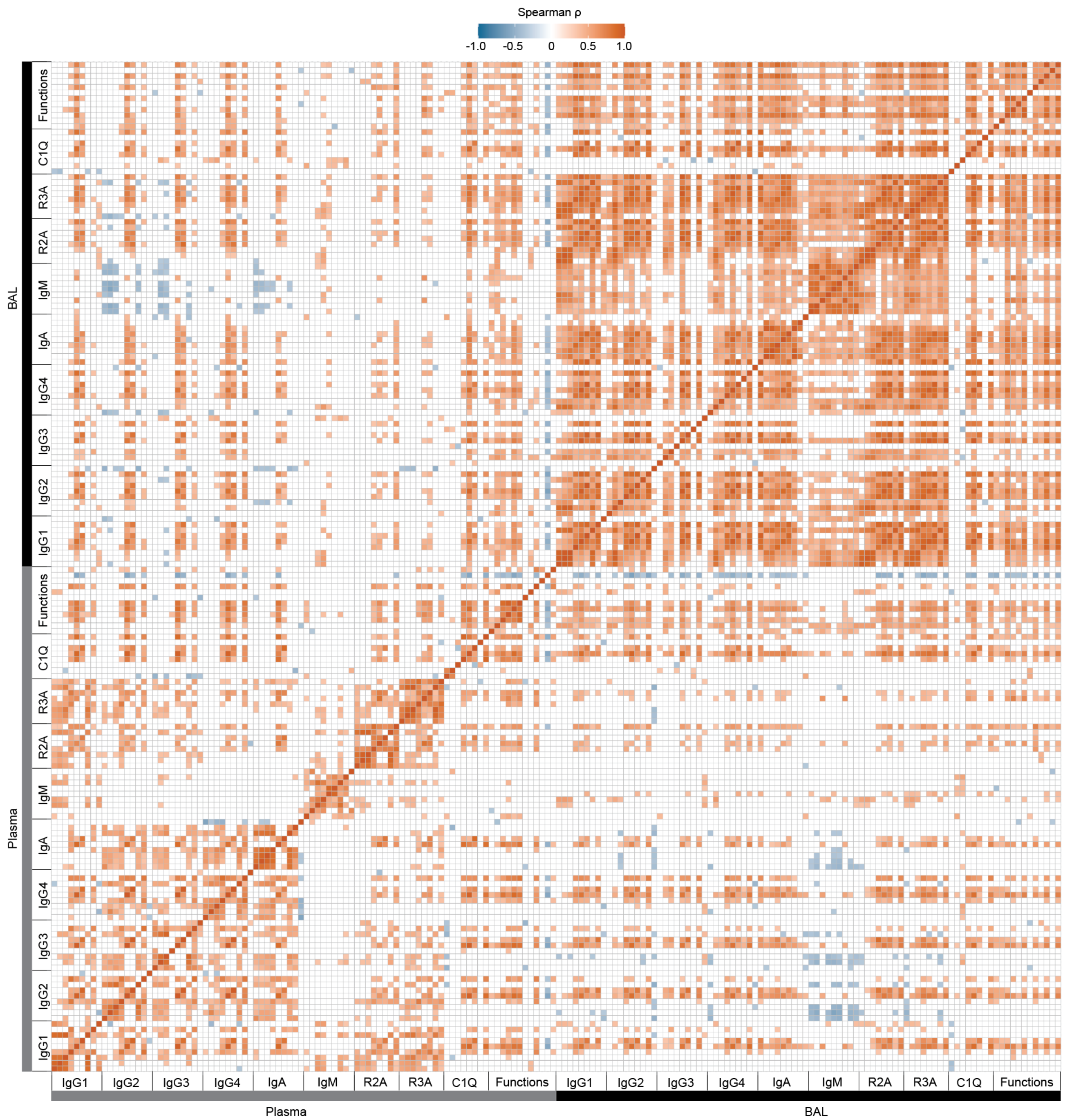


**Figure S5 | Correlation matrix of each antibody feature at peak immunogenicity.** Positive and negative correlations are colored orange and blue respectively. Spearman correlations with unadjusted p-value < 0 are white.
